# A tumor necrosis factor-α responsive cryptic promoter drives overexpression of the human endogenous retrovirus *ERVK-7*

**DOI:** 10.1101/2025.03.09.642286

**Authors:** Sojung Lee, Yin Yee Ho, Suyu Hao, Yingqi Ouyang, U Ling Liew, Ashish Goyal, Stephen Li, Jayne A. Barbour, Mu He, Yuanhua Huang, Jason W. H. Wong

**Affiliations:** School of Biomedical Sciences, The University of Hong Kong, Hong Kong SAR, China; Centre for Oncology and Immunology, Hong Kong Science Park, Hong Kong SAR, China; Cancer Epigenomics, German Cancer Research Center (DKFZ), Heidelberg, Germany; Center for Translational Stem Cell Biology, Hong Kong Science and Technology Park, Hong Kong SAR, China; Department of Statistics and Actuarial Science, The University of Hong Kong, Pokfulam, Hong Kong SAR, China

## Abstract

Endogenous retroviruses (ERVs) shape human genome functionality and influence disease pathogenesis, including cancer. *ERVK-7*, a significant ERV, acts as an immune modulator and prognostic marker in lung adenocarcinoma (LUAD). Although *ERVK-7* overexpression has been linked to the amplification of the 1q22 locus in approximately 10% of LUAD cases, it predominantly arises from alternative regulatory mechanisms. Our findings indicate that the canonical 5’ long terminal repeat (LTR) of *ERVK-7* is methylated and inactive, necessitating the use of alternative upstream promoters. We identified two novel transcripts, *ERVK-7.long* and *ERVK-7.short*, arising from distinct promoters located 2.8 kb and 13.8 kb upstream of the 5’LTR of *ERVK-7*, respectively. *ERVK-7.long* is predominantly overexpressed in LUAD. Through comprehensive epigenetic mapping and single-cell transcriptomics, we demonstrate that *ERVK-7.long* activation is predetermined by cell lineage, specifically in small airway epithelial cells (SAECs), where its promoter displays tumor-specific H3K4me3 modifications. Single-cell RNA sequencing further reveals a distinct enrichment of *ERVK-7.long* in LUAD tumor cells and alveolar type 2 epithelial cells, underscoring a cell-type-specific origin. Additionally, inflammatory signaling significantly influences *ERVK-7* expression; TNF-α enhances *ERVK-7.long*, while interferon signaling preferentially augments *ERVK-7.short* by differential recruitment of NF-κB/RELA and IRF to their respective promoters. This differential regulation clarifies the elevated *ERVK-7* expression in LUAD compared to lung squamous cell carcinoma (LUSC). Our study elucidates the complex regulatory mechanisms governing *ERVK-7* in LUAD and proposes these transcripts as potential biomarkers and therapeutic targets, offering new avenues to improve patient outcomes.

## Introduction

Transposable element (TE) derived sequences comprise nearly half of the human genome(1), with 8.3% associated with endogenous retroviruses (ERVs)(2). Due to their human-specific nature, human endogenous retroviruses (HERVs) garner significant interest among all TEs(3). In particular, HML2, a subgroup of the HERV-K family, has attracted substantial scientific attention due to its capacity for active transcription of RNA, coding for functional proteins and its association with various cancers(4). While most HERVs are silenced by DNA methylation or epigenetic alterations, in those that remain active, the promoter of HERVs is typically located in the 5’ long terminal repeat (LTR) region(5).

A recent study(6) has elucidated the role of HERV envelope glycoproteins in modulating responses of lung-resident B cells in lung adenocarcinoma (LUAD). The study pinpointed *ERVK-7* (also called HERV-K102 or HML2_1q22) on chromosome 1q22 as the HERV responsible, with *ERVK-7* specifically overexpressed in LUAD compared with lung squamous cell carcinoma (LUSC) or healthy controls(6). Importantly, the study demonstrated that antibody responses targeting the *ERVK-7* envelope glycoprotein increase following ICB treatment. Furthermore, elevated pre-treatment levels of *ERVK-7* were significantly associated with improved overall survival(6). These findings highlight the crucial clinical relevance of *ERVK-7* and emphasize the need to study mechanisms driving its upregulation. While chromosomal amplification has been proposed as a regulatory mechanism, it accounts for only ∼10% of lung adenocarcinoma (LUAD) cases(6). Therefore, a comprehensive investigation is necessary to elucidate the epigenetic regulation of *ERVK-7*.

Here, we present a comprehensive investigation of *ERVK-7* regulation. Rather than using its canonical promoter, which we found to be methylated and non-functional, *ERVK-7* employs two previously unidentified upstream transcription start sites. This novel regulatory mechanism generates distinct transcripts, *ERVK-7.short* and *ERVK-7.long*, each driven by specific transcription factors. Together, the regulation of these transcripts explains the variance in ERVK-7 expression among LUAD patients and its selective upregulation in LUAD compared with LUSC. Our findings provide the first mechanistic explanation for *ERVK-7*’s overexpression in LUAD.

## Results

### Identification of *ERVK-7* transcripts in Lung Adenocarcinoma

To accurately identify the TSS of *ERVK-7*, it is crucial to obtain the complete sequence of its transcripts. To do so, we obtained Pacific Biosciences (PacBio) Iso-Seq long-read RNA sequencing data for the LUAD cell line NCI-H1975(7). We have identified two distinct transcripts, designated as *ERVK-7.short* and *ERVK-7.long*, depicted in dark blue in Figure 1. These transcripts originate from uniquely positioned promoters: the promoter for *ERVK-7.long* is situated approximately 13.8 kb upstream, while the promoter for *ERVK-7.short* commences roughly 2.8 kb upstream of the gene annotated as *ERVK-7* in the RepeatMasker(8) database. These transcripts lead to the transcription of the complete *ERVK-7* gene, whereas other transcripts lead to only partial expression of *ERVK-7*, particularly missing the region encoding the *Env* protein. Interestingly, we did not detect any transcripts spliced to the ORF of *Env*, a splicing pattern typically associated with the production of the *Env* protein(9, 10).

**Figure 1:**
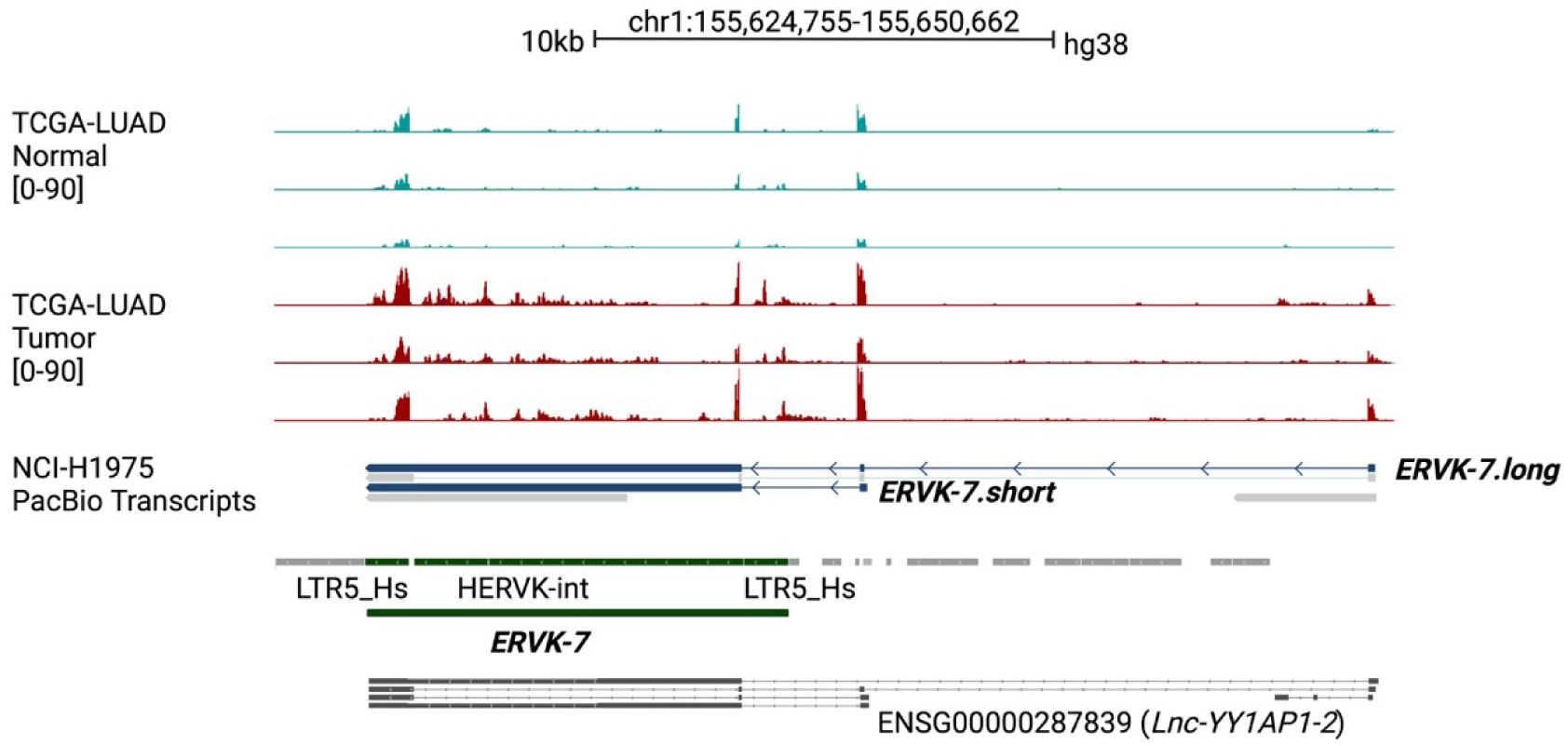
Identification of two *ERVK-7* transcripts from Pacbio Iso-seq data. The first six tracks display three pairs of TCGA LUAD normal (blue) and tumor (red) patient data (TCGA-44-2668, TCGA-38-4632, TCGA-44-6147) based on unique read coverage. The range for each track is indicated inside brackets. The NCI-H1975 (LUAD cell line) track presents transcripts identified from PacBio Iso-Seq data. The first blue-colored transcript represents *ERVK-7.long*, and the second blue-colored transcript represents *ERVK-7.short*. Subsequent tracks include RepeatMasker annotations; regions corresponding to *ERVK-7* are colored green. Two flanking LTR elements, LTR5Hs, span both the start and end of *ERVK-7*, with the internal part known as HERVK-int. *ERVK-7.short* and *ERVK-7.long* are indicated to originate from different promoters. *ERVK-7.long*, the longer transcript, begins approximately 13.8 kb from the original *ERVK-7* annotation. *ERVK-7.short*, the shorter transcript, starts approximately 2.8 kb from the original *ERVK-7* annotation. The final track (grey) displays transcripts from the V46 comprehensive dataset, labeled *Lnc-YY1AP1*.

We assessed three paired normal and tumor TCGA-Lung Adenocarcinoma (TCGA-LUAD) samples(11) (Fig. 1). Generally, *ERVK-7* expression was higher in tumor tissue compared to normal. In particular, the first exon of *ERVK-7.long* was significantly upregulated in tumor. A similar trend can be observed with the inclusion of multi-mapped reads (Supp. Fig. S1A).

To confirm that these transcripts indeed lead to the expression of *ERVK-7*, we identified spliced reads originating from the transcripts’ exons that entered the internal coding regions of *ERVK-7* (Supp. Fig. S1B). This suggests that the overexpression of the alternative *ERVK-7* transcripts may be linked to *ERVK-7* overexpression in LUAD (Fig. 1). The current GENCODE release (v47 hg38.p14) comprehensive dataset(12) contains six transcripts in the corresponding region, all labeled as lncRNA for the nearby upstream gene *YY1AP1* (Fig. 1A). As far as we are aware, no studies have been conducted on these transcripts.

### 5’ LTR of *ERVK-7* is epigenetically silenced with minimal evidence of transcription initiation across human cell types

The canonical promoter of full-length ERVs is located in the R region of the 5’ LTR(5) (Fig. 2A). Among the various subfamilies of LTR, *ERVK-7* is associated with LTR5HS, which is recognized for having the highest promoter activity compared to other subfamilies such as LTR5A or LTR5B. Previous studies have consistently shown that LTR5Hs typically exhibit the strongest promoter activity compared to LTR5A or LTR5B, and can effectively induce transcription in luciferase assays(5). However, our analysis of 114 primary cell types from ENCODE(12) revealed that only 11 had junction reads spanning both the 5’ LTR and coding region (HERVK-int) of *ERVK-7*. Moreover, the frequency of these captured junction reads was shallow, with counts per million (CPM) averaging approximately 0.05 (Fig. 2B). Thus, we investigated potential reasons for the 5’ LTR region, LTR5Hs of *ERVK-7*, to be mostly non-functional as a promoter.

**Figure 2:**
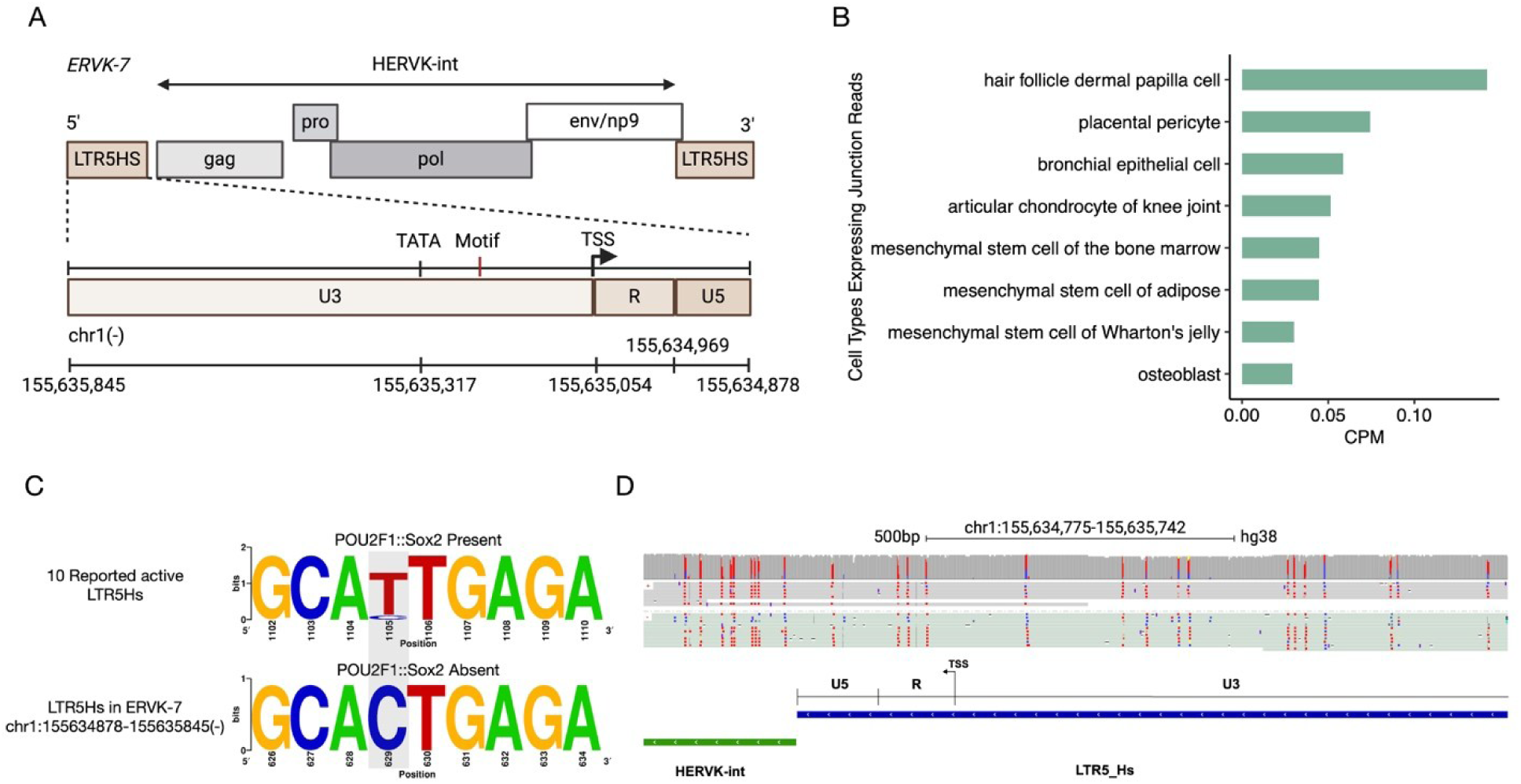
Promoter in 5’ LTR5Hs of *ERVK-7* is largely silent. (A) Schematic diagram showing putative promoter for *ERVK-7* in the 5’ LTR5Hs. (B) Junction reads covering LTR5Hs and HERVK-int for 57 primary cells types from ENCODE. (C) Motif graph showing a T > C substitution (Grey highlited aread) when comparing in 10 reported active LTR5Hs with the *ERVK-7* 5’ LTR5Hs. (D) A549 Nanopore direct DNA-sequencing data showing methylation status over a portion of *ERVK-7* 5’ LTR5Hs. Unmethylated CpG sites (Blue) and methylated CpG sites (Red).

We first compared genomic sequence differences in LTR5Hs with 10 LTR5Hs that were previously found to have transcriptional activity(13) (Supp. Table S1). We identified a T>C single nucleotide variant (motif label in Fig. 2A, Grey area in Fig. 2C), potentially leading to a change in the binding of POUS2F1::SOX2 (Fig. 2C), which has been shown to be an important transcription factor regulating the expression of HERV-K(14).

We further hypothesized that DNA methylation might be a major driver of inactivity. As the LTR5Hs region is highly repetitive, the mappability of whole-genome bisulfite sequencing (WGBS) data is limited. Thus, we generated Nanopore direct DNA-sequencing data from A549 cells and quantified DNA methylation at the 5’ LTR5Hs of *ERVK-7*. Across R and U5 regions, most of the CpG sites were methylated (Fig. 2D). This finding confirms that the 5’ LTR5Hs of *ERVK-7* is silenced, but the hijacking of alternative upstream transcript start sites and exons likely rescues the expression of *ERVK-7*.

### *ERVK-7.long* expression underlies the upregulation of *ERVK-7* in LUAD and LUSC samples

Frequent amplification of chromosome 1q22 observed was described as a factor contributing to the overexpression of *ERVK-7* in LUAD patients. However, only around 10% of LUAD patient genomes have this amplification (Supp. Fig. S2A). We hypothesized that the two novel transcripts associated with *ERVK-7*, designated as *ERVK-7.short* and *ERVK-7.long*, may drive its expression. Given the substantial overlap between these transcripts, we first defined a unique 111 bp region specific to *ERVK-7.short* to facilitate accurate quantification (Fig. 3A, Supp. Table S2). For *ERVK-7.long*, we utilized the distal exon unique to this transcript, while for *ERVK-7* itself, we employed the original region as defined by RepeatMasker (Fig. 3A). This approach enabled us to distinguish and quantify the expression of each transcript independently without the need to deconvolute the expression of individual *ERVK-7* transcripts.

**Figure 3:**
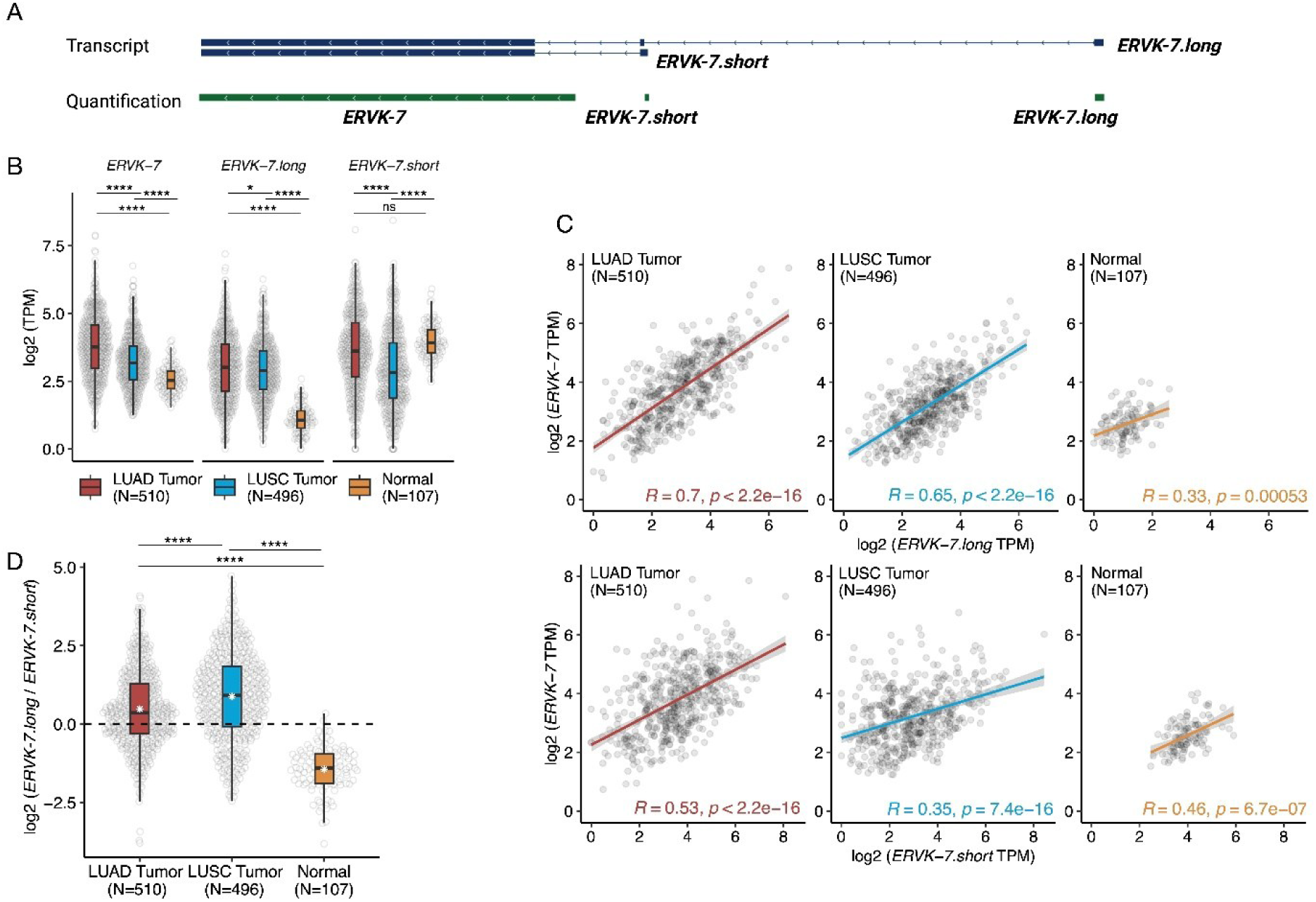
Expression of *ERVK-7* in TCGA lung cancer cohort study. (A) Track showing original transcripts for *ERVK-7.long* and *ERVK-7.short.* The green track represents the GTF file used for quantification for *ERVK-7* transcripts. For *ERVK-7.short* and *ERVK-7.long*, quantified regions are unique to the respective transcripts. *ERVK-7* was retrieved from RepeatMasker annotation. (B) Plot showing quantified log2 TPM values for three *ERVK-7* transcripts. T-test performed (C) Top panel: Spearman’s rank correlation test was conducted between log2 TPM of *ERVK-7.long* and log2 TPM of *ERVK-7*. Bottom panel: Spearman’s rank correlation test was conducted between the log2 TPM of *ERVK-7.short* and the log2 TPM of *ERVK-7*. Red: LUAD tumor samples; Blue: LUSC tumor samples; Yellow: LUAD and LUSC normal samples. (D) log2 Ratio of *ERVK-7.long* and *ERVK-7.short* in TCGA LUAD tumor (Red), LUSC tumor (Blue), and Normal (Yellow). White start in boxplot represents mean of each group. See method for detail. Student t-test conducted. (NS: p > 0.05, ∗: p ≤ 0.05, ∗∗: p ≤ 0.01, ∗∗∗: p ≤ 0.001, ∗∗∗∗: p ≤ 0.0001).

We investigated the expression patterns of the *ERVK-7* transcripts in TCGA LUAD and LUSC cohorts. *ERVK-7.long* and *ERVK-7* showed significant upregulation in tumors compared with normal tissue (mean (*ERVK-7.long* TPM) = 1.29; mean (*ERVK-7* TPM) = 5.37), specifically with LUAD patients (mean (*ERVK-7.long* TPM) = 10.77; mean (*ERVK-7* TPM) = 18.47) exhibiting higher expression of both transcripts compared with LUSC patients (mean (*ERVK-7.long* TPM) = 9.06; mean (*ERVK-7* TPM) = 10.86) (Fig. 3B; Student t-test, p-value (*ERVK-7.long* LUAD vs LUSC) = 0.016, p-value (*ERVK-7.long* LUAD vs Normal) < 2.2e-16, p-value (*ERVK-7* LUAD vs LUSC) = 3.01e-12, p-value (*ERVK-7.* LUAD vs Normal) < 2.2e-16). *ERVK-7.short* displayed a different trend, with normal generally having higher expression than tumour (Fig. 3B; mean (LUAD) = 18.74, mean (LUSC) = 12.49, mean (Normal) = 16.46, Student t-test, p-value (*ERVK-7.short* LUAD vs LUSC) = 9.4e-06, p-value (*ERVK-7.short* LUAD vs Normal) = 0.08).

To determine whether *ERVK-7.long* or *ERVK-7.short* is the primary contributor to *ERVK-7* expression in TCGA-LUAD and LUSC, we conducted a series of correlation tests. *ERVK-7.long* and *ERVK-7* were generally more correlated (LUAD: R=0.7; LUSC: R=0.65) (Fig. 3C) compared with *ERVK-7.short* and *ERVK-7* (LUAD: R=0.53; LUSC: R=0.35) (Fig. 3C) in both tumor subtypes. This suggests that *ERVK-7.long* is the primary contributor to *ERVK-7* upregulation in tumor samples. We further quantified TCGA samples using the same region from Ng et al. (trans996c17b9912b02e8)(6) that encodes the *ERVK-7 Env* protein and found a similar trend (Supp. Fig. S2B-C). To further ensure that our observation is independent of 1q22 amplification, we selected the subset of TCGA lung cohort patients (LUAD: N=462, LUSC: N=467, Normal: N=98) without this amplification. *ERVK-7.long* remains the primary driver of *ERVK-7* upregulation (Supp. Fig. S3A-B).

Finally, we generated a regression model (see Methods) to assess the relative contribution of *ERVK-7.short* and *ERVK-7.long* to *ERVK-7* in each sample. Based on this model, samples with greater contribution from *ERVK-7.short* compared with *ERVK-7.long* will have a log ratio < 0, whereas those with greater contribution from *ERVK-7.long* will be > 0. It is evident that for both LUAD and LUSC samples, *ERVK-7.long* is the major contributor to *ERVK-7* expression, whereas in normal lung, *ERVK-7.short* is the primary driver (mean (LUAD) = 0.48, mean (LUSC) = 0.88, mean (Normal) = −1.44, Fig. 3D). This suggests that *ERVK-7.long* upregulation is critical in triggering *ERVK-7* upregulation in lung cancer.

### Cell of origin determines the expression of the *ERVK-7.long* transcript

Given the contribution of *ERVK-7.long* in driving overexpression in LUAD, we sought to determine the mechanisms underlying its apparent specificity in lung cancers. We comprehensively analyzed available epigenomic data to investigate whether epigenetic alterations between normal and tumor might contribute to this tissue-specific expression pattern. To examine chromatin accessibility, we first compared ATAC-seq data from normal lung tissue and A549 cells, which showed consistent profiles (Fig. 4A).

**Figure 4:**
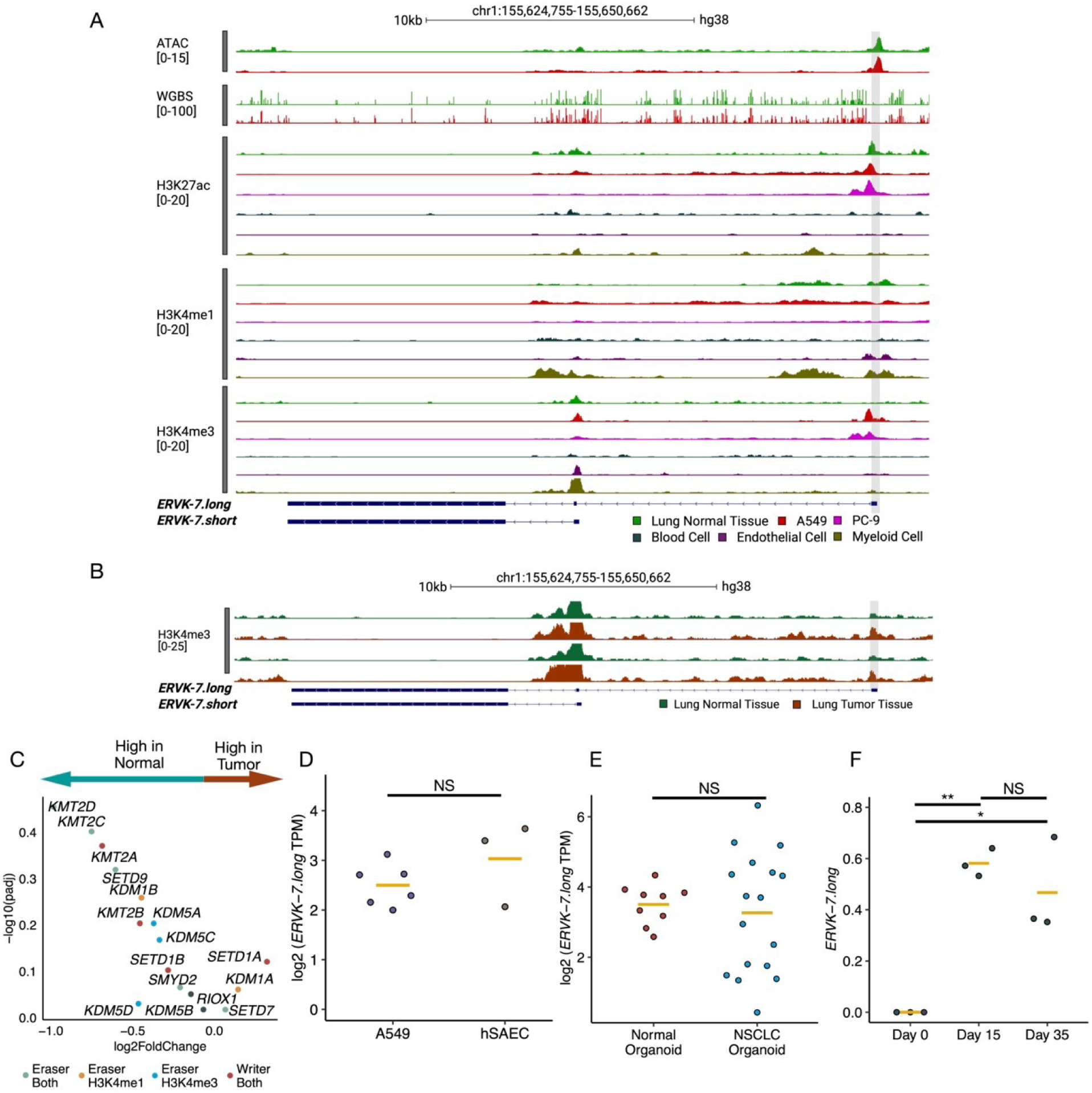
Cell type specificity of *ERVK-7.long* promoter and transcript. (A) Analysis includes ATAC and WGBS for A549 (red) and normal lung (green). Displayed are three epigenetic tracks for normal lung (green), A549 (red), PC-9 (pink), blood mononuclear cells (black), endothelial cells (brown), and myeloid cells (khaki), featuring H3K27ac, H3K4me1, and H3K4me3. The grey highlight marks the promoter region of *ERVK-7.long*. Data were sourced from ENCODE. The number inside brackets on each track indicate the following: ATAC shows the fold change over control; WGBS reveals the methylation state at the CpG minus strand; H3K27ac, H3K4me1, and H3K4me3 depict the fold change over control. (B) Two datasets of H3K4me3 ChIP-seq compare normal lung (green) and lung tumor (red) tissues. The grey highlighted area indicates the promoter of *ERVK-7.long*. 25 fold change over the control is applied as a peak threshold. (C) Differential expression analysis performed between TCGA LUAD tumor vs normal. Volcano plot showing histone K4 readers and writers genes. Light green: Eraser genes for both H3K4me3 and H3K4me1. Orange: eraser genes for H3K4me1. Blue: eraser genes for H3K4me3. Red: writer for both H3K4me3 and H3K4me1. (D) *ERVK-7.long* log2 TPM values between A549 (Dark purple) and HSAEC (Brown). Student t-test conducted. Yellow line represents mean value for each group. (E) *ERVK-7.long* log2 TPM values between normal organoid (Red) and tumor organoid (Blue). Student t-test conducted. Yellow line represents mean value for each group. (F) *ERVK-7.long* log2 TPM values among PSCs with three time points (Day 0, Day 15, and Day 35). Student t-test conducted. Yellow line represents mean value for each group. (NS: p > 0.05, ∗: p ≤ 0.05, ∗∗: p ≤ 0.01, ∗∗∗: p ≤ 0.001, ∗∗∗∗: p ≤ 0.0001).

We then examined H3K27ac, H3K4me1, and H3K4me3 ChIP-sequencing (ChIP-seq) profiles for normal lung tissue, two LUAD cell lines (A549, PC-9) and three primary cell types: endothelial cells of umbilical vein, myeloid progenitor (CD34-positive) cells, and peripheral blood mononuclear cells from the ENCODE project(15) (Fig. 4A). WGBS and H3K27ac data indicated that the *ERVK-7.long* promoter is in open chromatin and an active region in both LUAD cell lines and normal lung tissue (Fig. 4A). However, LUAD cell lines exhibited clear evidence of H3K4me3 deposition compared to all other samples, suggesting that this region acts as a promoter specifically in LUAD tumor cells (Fig. 4A). For further verification, deposition of H3K4me3 in the promoter of *ERVK-7.long* was observed across five additional NSCLC cell lines (Supp. Fig. S4A). In endothelial cells, myeloid cells, and normal lung tissue, the presence of H3K4me1 together with H3K27ac suggested that it functions as an enhancer (Fig. 4A). Examination of the ENCODE Registry of candidate cis-Regulatory Elements (cCRE) database further confirmed that the *ERVK-7.long* promoter acts as an enhancer in most cell types(16).

To ensure that our observation is not biased by comparing cell lines against tissue, we further compared H3K4me3 levels between normal and tumor lung tissue(17). Indeed, even in lung tumour tissue, there is an evident increase in H3K4me3 signal at the promoter of *ERVK-7.long* compared with normal lung tissue (Fig. 4B, Supp. Fig. S4B). In bulk tissue, H3K4me3 signal at the promoter of *ERVK-7.long* is still substantially lower than *ERVK-7.short* as *ERVK-7.short* is broadly active across most cells types while *ERVK-7.long* is cell type specific.

This observation suggests that H3K4me3 deposition may convert the enhancer region into a promoter, resulting in the transcription of *ERVK-7.long*. We examined the expression of known H3K4 methylation eraser and writer genes in paired normal and tumor samples from TCGA LUAD and LUSC cohorts. However, these genes did not show significant differential expression (cutoff: adjusted p-value < 0.05; −log10 (adjusted p-value) > 1.30) (Fig. 4C, Supp. Fig. S4C).

An alternative explanation is that *ERVK-7.long* lung tumor cells may have inherited the epigenetic state from a smaller originating population of normal lung epithelial progenitor cells. Non-small cell lung cancer is known to derive from small airway epithelial cells (SAEC)(18, 19). A cosine similarity test of the entire transcriptome revealed high similarities between these cell lines (Supp. Fig. S4C). We measured *ERVK-7.long* expression in human SAEC (hSAEC) and found similar levels in hSAEC and A549(19) (Fig. 4D). We confirmed these findings using tumor organoids derived from NSCLC tumor cells(20, 21) and normal organoids from normal lung epithelial cells(22, 23). Both tumor and normal organoid showed similar *ERVK-7.long* expression (Fig. 4E; mean (normal organoid) = 3.5; mean (tumor organoid) = 3.27; Student t-test, p-value = 0.6042). These results suggest that the expression of *ERVK-7.long* is predetermined by its cell type of origin. hSAEC already possess H3K4me3 deposition (Supp. Fig. S5A)(24), with *ERVK-7.long* expression. Moreover, during the differentiation of human pluripotent stem cells (PSCs) into lung epithelial cell progenitors at days 0, 15, and 35(25), activation of the *ERVK-7.long* promoter was observed from day 15, coinciding with the transition to lung progenitors (Fig. 4F, Supp. Fig. S5B; TPM mean (Day 0) = 0, mean (Day 15) = 0.58, mean (Day 35) = 0.47, p-value (Day 0 vs Day 15) = 0.003, p-value (Day 0 vs Day 35) = 0.04, p-value (Day 15 vs Day 35) = 0.404, Student t-test). These results support the hypothesis that *ERVK-7.long* expression is intrinsically linked to lung cell lineage specification with the population expanded in lung cancer cells.

### Single-cell RNA-seq analysis confirms *ERVK-7.long* is expressed in a cell type-specific manner

To validate our findings, we analyzed scRNA-seq from lung tumor patients. We chose a lung scRNA-seq dataset generated using SMART-seq2(26) as its full-length transcript coverage ensures the ability to differentiate the expression of *ERVK-7.short* and *ERVK-7.long* at the single-cell level. This dataset included 26,485 cells from 49 samples (45 LUAD, one LUSC, and three adjacent normal tissue samples) covering primary tumors and metastases(27). Cells were initially categorized into six subtypes: endothelial, fibroblast, melanocytes, epithelial, immune, and tumor cells.

Our analysis showed that *ERVK-7.long* expression is highly specific to certain cell types, while *ERVK-7.short* and *ERVK-7* are expressed across all cell types (Fig. 5A). This pattern was consistent in t-SNE plots for primary LUAD patients (Supp. Fig. S6). Among cell types expressing *ERVK-7.long*, tumors showed the highest proportion, followed by fibroblasts and epithelial cells (Fig. 5B). For *ERVK-7.short* and *ERVK-7*, tumor and immune cells showed similar expression proportions (Fig. 5B).

**Figure 5:**
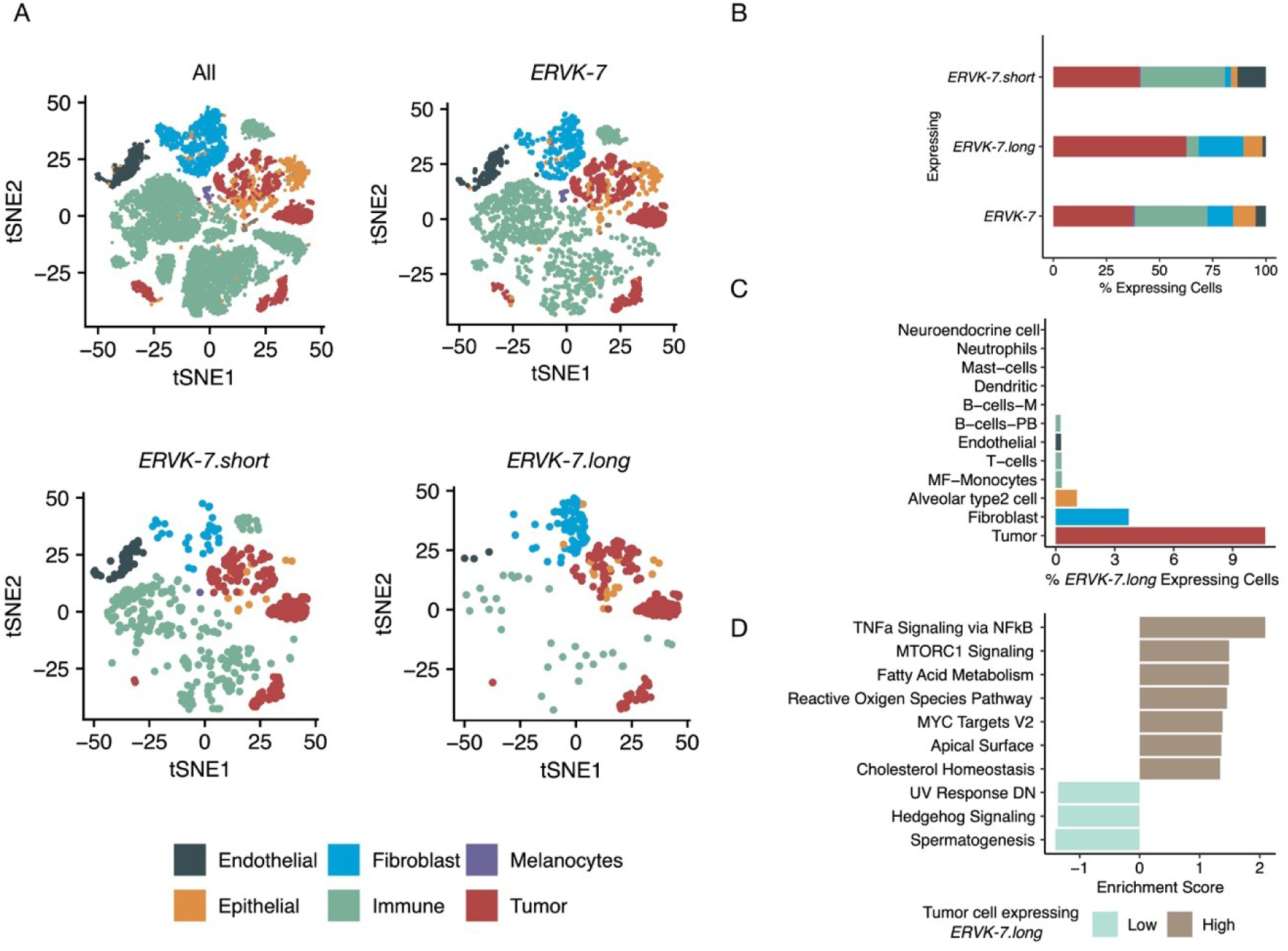
scRNA supports *ERVK-7.long* expression in specific cell types. (A) t-distributed stochastic neighbor embedding (t-SNE) plot of SMART-seq2 scRNA-seq data from lung cancer and normal samples. Subpanels show all cells or cells expressing specific *ERVK-7* transcripts. Six board cell types marked: Dark grey: Endothelial, Blue: Fibroblast, Purple: Melanocytes, Orange: Epithelial, Green: Immune, Red: Tumor. (B) Cell types proportions for three *ERVK-7* related transcripts. % of *ERVK-7.long* expressing cell types: tumor (62.7%), fibroblast (21.1%), epithelial (8.9%), immune (5.7%), endothelial (1.6%). % of *ERVK-7.short* expressing cell types: tumor (40.6%), fibroblast (2.97%), epithelial (2.97%), immune (39.6%), endothelial (13.4%). % of *ERVK-7* expressing cell types: tumor (37.4%), fibroblast (12.1%), epithelial (10.5%), immune (34.2%), endothelial (4.8%). (C) Proportion of cells expressing *ERVK-7* in each detail cell annotation. % of *ERVK-7.long* expressing cell types: tumor (10.7%), fibroblast (3.72%), AT2 (1.08%), MF-Monocytes (0.3%), T-cells (0.3%), endothelial (0.3%), B-cells-PB (0.25%), B-cells-M (0%), Dendritic (0%), Mast-cells (0%), Neutrophils (0%), Neuroendocrine cell (0%). Only cell types with having more than 600 cells are chosen. (D) Top 10 most significant MSigDB Hallmark pathways based on GSEA comparing *ERVK-7.long* lowly or non-expressing tumor cells (Turquoise) and *ERVK-7*.long highly expressing tumor cells (Brown).

Cell type-specific analysis revealed a distinct pattern of *ERVK-7.long* expression, with tumor cells exhibiting the highest levels, followed by fibroblasts and Alveolar Type 2 (AT2) cells (Fig. 5C). The presence of *ERVK-7.long* expression in AT2 cells aligns with expectations, given that AT2 cells constitute a significant fraction of SAEC(28). Notably, AT2 and tumor cells clustered together, demonstrating high transcriptomic similarity (Fig. 5A). This clustering aligns with the established understanding that LUAD predominantly originates from AT2 cells(29–31), with scRNA-seq study showing that these share transcriptomic similarities(32). Moreover, a recent study demonstrated that malignant AT2-like cells in LUAD exhibit progressive chromosomal alterations across cancer stages, providing strong evidence for AT2 cells as the likely cellular origin of this malignancy(32). Given this lineage relationship, the elevated *ERVK-7.long* expression in tumor cells likely stems from their AT2 cell origin.

While the proportion of cell types may vary depending on several factors, including sample preparation, we observed a significantly higher percentage of *ERVK-7.long*-expressing cell types (AT2, fibroblast, and tumor) per patient in primary tumors compared to normal tissue (Supp. Fig. S6B; mean (tumor) = 41.14%, mean (normal) = 25.5%; Student’s t-test, p = 0.04). This observation demonstrates an expansion of *ERVK-7.long*-expressing cells in tumors relative to normal tissue, contributing to its expression detected in LUAD bulk RNA-seq data.

Notably, neuroendocrine cells showed no detectable *ERVK-7.long* expression (Fig. 5C). Given that small cell lung cancer (SCLC) primarily arises from neuroendocrine cells(33, 34), we examined *ERVK-7* transcript levels in SCLC cell lines(35). Compared to the A549 line, all six SCLC cell lines lacked *ERVK-7.long* expression, correlating with reduced overall *ERVK-7* expression (Supp. Fig. S7). These results confirm that LUAD tumour cell’s ability to express *ERVK-7.long* expression is likely inherited from its cell type of origin, the AT2 epithelial cells.

While our data established cellular origin as a key determinant of *ERVK-7.long* expression, the high variance observed in *ERVK-7.long* levels among LUAD samples remained unexplained. To address this question, we performed Gene Set Enrichment Analysis (GSEA) comparing tumor cells with high versus low *ERVK-7.long* expression (see Methods). Notably, the most significantly upregulated pathway identified was TNF-α signaling via the NF-κB pathway (Fig. 5D). This finding suggests a potential role for TNF-α signaling or NF-κB regulating the expression of *ERVK-7.long*.

### TNF-α Signaling via NF-κB/RELA Activation Regulate *ERVK-7.long* Expression in Lung Adenocarcinoma

To further investigate the role of TNF-α signaling, we examined transcription factor (TF) binding to the *ERVK-7.long* promoter. Screening with cistromeDB revealed RELA, a subunit of nuclear factor NF-κB, as one of the most highly enriched TFs at this locus (Supp. Fig. S8A). This is consistent with the scRNA-seq analysis linking TNF-α signaling and NF-κB with *ERVK-7.long* expression.

To validate our observations, we analyzed ChIP-seq data for RELA in the A549 cell line(36). This confirmed RELA binding to the *ERVK-7.long* promoter (Fig. 6A). Further examination of A549 cells treated with TNF-α(36, 37), revealed that TNF-α treatment alone significantly upregulated both *ERVK-7* (Paired t-test, p = 0.009) and *ERVK-7.long* expression (Paired t-test, p = 0.02), while *ERVK-7.short* remained largely unaffected (Paired t-test, p = 0.73; Fig. 6B). We subsequently quantified three *ERVK-7* transcripts in the 11-18 LUAD cell line treated transduced with IκB-α, a natural inhibitor of NF-κB, or RelAS536E, a constitutively active RELA mutant(38). Both *ERVK-7.long* and *ERVK-7* exhibited significant increases in response to RelAS536E (Student’s t-test, p = 0.002 and p = 0.013, respectively) and marked decreases following IκB-α treatment (Fig. 6C; Student’s t-test, p = 0.004 and p = 0.043, respectively). *ERVK-7.short* displayed a similar trend between control and RelAS536E conditions (Fig. 6C; Student’s t-test, p = 0.04); however, its decrease with IκB-α treatment did not reach statistical significance (Fig. 6C; Student’s t-test, p = 0.053). To further validate NF-κB’s role in A549, we examined the effect of two NF-κB inhibitors, PA and PS1145, on TNF-α-induced A549 cells(36). Treatment with these inhibitors significantly reduced *ERVK-7.long* and *ERVK-7* expression to control levels or lower, underscoring NF-κB’s crucial role in regulating *ERVK-7.long* and its function as an upstream regulator of *ERVK-7* (Supp. Fig. S8B, Supp. Table S3).

**Figure 6:**
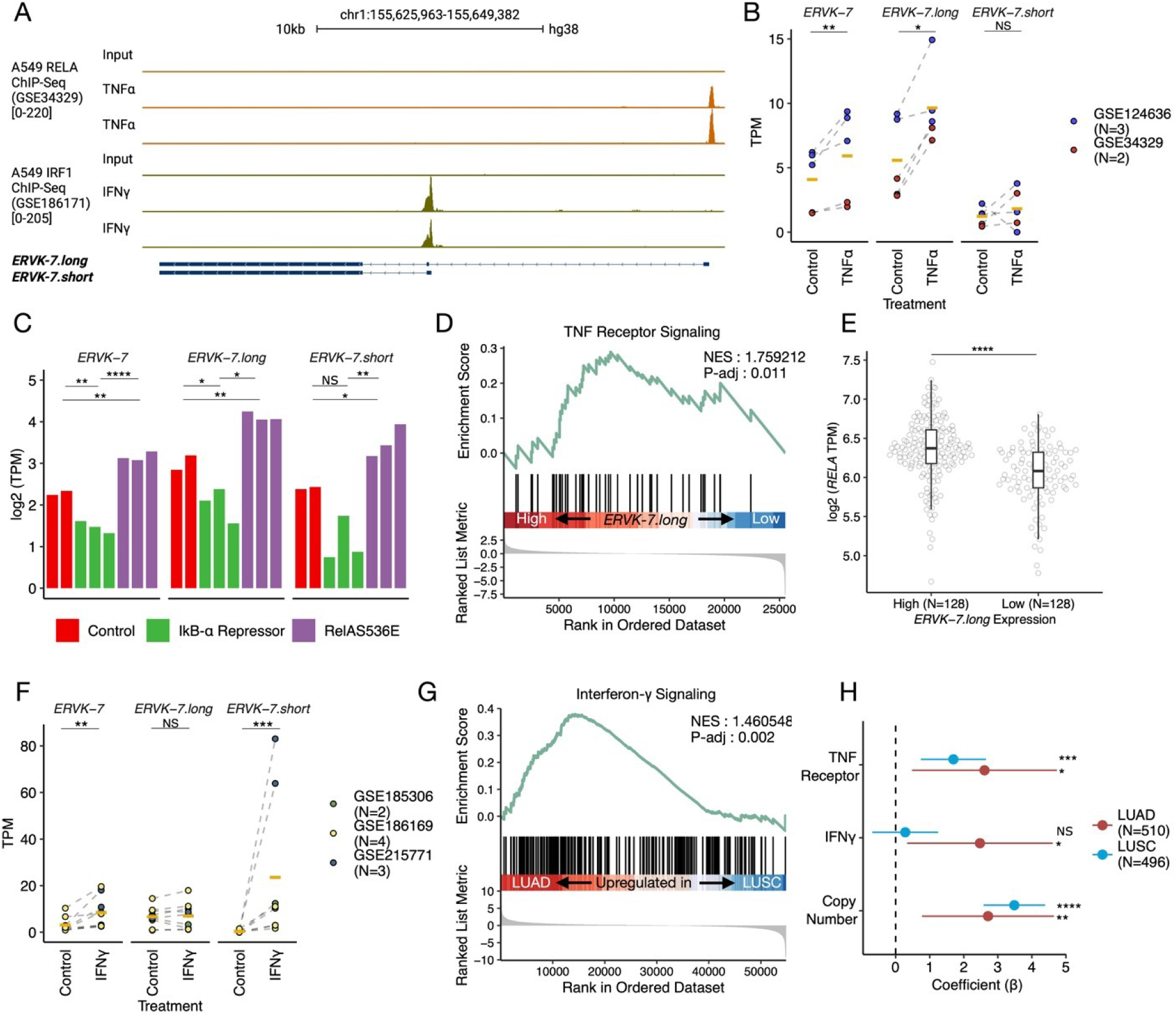
Pathways Driving *ERVK-7* Upregulation in Lung Adenocarcinoma. (A) RELA ChIP-seq with two replicates of TNF-α-induced A549 (orange) and IRF1 ChIP-seq with two replicates for IFN-γ-induced A549 (green). The input represents A549 DNA not subject to immunoprecipitation from the respective experiments. The range for each track is indicated inside brackets. (B) Log2 TPM values for three *ERVK-7* transcripts. Dot colors represent individual studies with GEO numbers labeled for each dataset. Dashed lines connect samples paired by study. Conditions include untreated A549 (control) and TNF-α-induced A549 (TNFα). Yellow lines indicate the mean of each group. A paired t-test was applied. (C) Log2 TPM values for three *ERVK-7* transcripts across three conditions: control A549 (red), A549 expressing the IκB-α repressor (green), and A549 expressing RelA S536E (purple). IκB-α is known as a repressor of the NF-kB pathway, and RelA S536E as an activator of RelA. Student’s t-tests were performed. (D) GSEA plot for TNF receptor signaling between TCGA-LUAD samples with high *ERVK-7.long* (n=128) and low *ERVK-7.long* (n=128) expressions. (E) RELA expression in Log2 TPM for high *ERVK-7.long* (n=128) and low *ERVK-7.long* (n=128) expressing groups. Student’s t-tests were performed. (F) Log2 TPM values for three *ERVK-7* transcripts Dot colors represent individual studies with GEO numbers labeled for each dataset. Dashed lines connect samples paired by study. Conditions include untreated lung cancer cell lines (control) and IFN-γ-induced lung cancer cell lines (IFNγ). Yellow lines indicate the mean of each group. A paired t-test was applied. (G) GSEA plot for two interferon-γ pathways between TCGA-LUAD and LUSC tumor samples. (H) Multivariable regression model for TCGA-LUAD (red) and LUSC (blue), with significance denoted as NS (p > 0.05), * (p ≤ 0.05), ** (p ≤ 0.01), *** (p ≤ 0.001), and **** (p ≤ 0.0001).

To investigate whether TNF-α also drives *ERVK-7* expression in clinical samples, we stratified TCGA-LUAD tumors based on *ERVK-7.long* expression levels. We selected the top 25% (N=128) and bottom 25% (N=128) of samples for analysis (Supp. Fig. S8C). Gene Set Enrichment Analysis (GSEA) of these groups revealed significant enrichment of TNF-α signaling via NFκB in samples with high ERVK-7.long expression (Supp. Fig. S8D, Normalized Enrichment Score [NES] = 1.24, adjusted p-value = 0.03). Given that the NFκB pathway can be activated by multiple signaling cascades(39, 40), we sought to investigate upstream of NFκB activation by examining TNF receptor signaling. GSEA also revealed significant enrichment of the TNF receptor signaling pathway in samples with high *ERVK-7.long* expression compared to those with low expression (Fig. 6D, NES = 1.76, adjusted p-value = 0.011). This enrichment was independent of 1q22 status, as we observed similar results in TCGA-LUAD tumor samples without 1q22 amplification (Supp. Fig. S8E-F, NES = 2.12, adjusted p-value = 0.0004). Moreover, *RELA* expression was significantly elevated in the high *ERVK-7.long* expression group (Fig. 6E; mean (High) = 6.37, mean (Low) = 6.03, Student’s t-test, p = 4.66e-10).

These results demonstrate that TNF-α signaling, acting through NF-κB and its subunit RELA, significantly upregulates *ERVK-7.long* expression in lung adenocarcinoma (LUAD). The observed upregulation of the TNF receptor signaling pathway further corroborates the activation of TNF-α signaling. Upon TNF-α induction, RELA binding to the *ERVK-7.long* promoter intensifies. Moreover, high *ERVK-7.long* expression in patient samples correlates with enriched TNF-α signaling and elevated RELA levels in LUAD.

### IFN-γ Drives Further Upregulation of *ERVK-7* in Lung Adenocarcinoma via *ERVK-7.short*

Despite significant upregulation of *RELA* expression in the *ERVK-7.long* high group, comparison of *RELA* expression among LUAD, LUSC, and normal samples revealed no significant changes or even higher expression in normal tissue (Supp. Fig. S9A; mean (LUAD tumor) = 6.22, mean (LUSC tumor) = 6.27, mean (Normal) = 6.29; Student’s t-test, p-value (LUAD tumor vs Normal) = 0.09034, p-value (LUSC tumor vs Normal) = 0.5874). Furthermore, the contribution to the expression of *ERVK-7.long* towards *ERVK-7* expression was even higher in LUSC vs LUAD (mean (LUSC tumor) = 0.88 vs mean (LUAD tumor) = 0.48, p-value (LUAD tumor vs LUSC tumor) = 1.5e-06, Student’s t-test, Fig. 3D, Supp. Table S3), consequently, we hypothesized that differential *ERVK-7.short* expression contributes to the disparity in *ERVK-7* expression between the two lung cancer subtypes.

The TSS of *ERVK-7.short* coincides with a previously described enhancer for *ERVK-7*(41). This region overlaps with a solo LTR element, *LTR12F*, which harbors an IRF binding site sensitive to IFN-γ stimulation. Our analysis suggests that *LTR12F* functions not only as an enhancer for *ERVK-7* but also directly as a promoter for the *ERVK-7.short* transcript, providing an alternative mechanism for driving *ERVK-7* expression.

To validate this hypothesis, we first confirmed interferon 1 (IRF1) binding using ChIP-seq data from A549(35) (Fig. 6A). Furthermore, analysis of three RNA-seq datasets from LUAD cell lines treated with IFN-γ(35, 42, 43) consistently demonstrated a significant increase in *ERVK-7.short* expression (Paired t-test, p = 0.0007). *ERVK-7* also showed increased expression (Paired t-test, p = 0.002), while *ERVK-7.long* remained unaffected (Paired t-test, p = 0.81; Fig. 6F). Notably, treatment of A549 cells with an IFN-γ inhibitor (AX0085) reversed or reduced *ERVK-7.short* and *ERVK-7* expression to levels comparable with A549 control (Supp. Fig. S9B, Supp. Table S3).

To confirm whether interferon signaling underlies the differential expression between LUAD and LUSC patient samples, we performed GSEA on the two subtypes. This analysis revealed significant enrichment of the IFN-γ pathway in LUAD compared to LUSC (Fig. 6G; NES = 1.46, adjusted p-value = 0.002). Thus, *ERVK-7.short* additionally contributes to *ERVK-7* expression in LUAD via elevated IFN-γ signaling.

To consolidate our findings, we constructed a multivariable regression model incorporating the key factors contributing to *ERVK-7* overexpression in both LUSC and LUAD tumor samples. Copy number variation was a significant determinant of *ERVK-7* expression in both cancer subtypes (Multivariate regression model, coefficient (LUAD) = 2.64, coefficient (LUSC) = 3.72, p-value (LUAD) = 0.00684, p-value (LUSC) = 3.56e-15). For TNF and IFN signaling, we observed distinct regulatory mechanisms in LUAD and LUSC. TNF receptor signaling pathway was significantly upregulated *ERVK-7.long* in both lung cancer types (Multivariate regression model, coefficient (LUAD) = 2.67, coefficient (LUSC) = 1.69, p-value (LUAD) = 0.013, p-value (LUSC) = 0.0005), while IFN-γ signaling was preferentially induced *ERVK-7.short* expression in LUAD (Fig. 6H; Multivariate regression model, coefficient (LUAD) = 2.64, coefficient (LUSC) = 0.25, p-value (LUAD) = 0.03, p-value (LUSC) = 0.61). This model accounts for *ERVK-7* expression patterns observed in LUAD and LUSC tumors, as well as in normal lung tissue, providing mechanistic insights into the complex regulatory network that controls *ERVK-7* expression in lung cancer (Supp. Fig. S10).

## Discussion

Our study reveals that 5’ LTR5Hs of *ERVK-7* is largely inactive due to methylation and SNVs. Consequently, upstream exons act as promoters, generating two distinct isoforms of *ERVK-7*: *ERVK-7.long* and *ERVK-7.short*. These transcripts are regulated by distinct transcription factors: RELA/NF-κB modulates *ERVK-7.long*, while IRF1 regulates *ERVK-7.short*. We used a regression model to show that *ERVK-7.long* is the major contributor to *ERVK-7* expression in lung cancer. Interestingly, while the *ERVK-7.long* promoter region typically shows enhancer characteristics across many cell types, specific lung epithelial cells such as AT2 display H3K4me3 deposition, leading to *ERVK-7.long* transcription.

Our findings suggest that cell type of origin predetermines *ERVK-7.long* activation and expression levels, with significant upregulation driven by TNF-α via NF-κB and IFN-γ pathways. Previous literature supports our observations, demonstrating HERV-K element upregulation in response to TNF-α in various diseases, including cancers, HIV infection, rheumatoid arthritis, and schizophrenia(44, 45).

*ERVK-7* upregulation is associated with a positive ICB response and a favorable prognosis(6). Our data suggests that TNF-α could enhance NF-κB pathways, acting as a pro-immunogenic factor in ICB context. However, the role of TNF-α via NF-κB in modulating ICB response remains controversial(46–48), as these pathways exhibit cell type-dependent functions. Our GSEA reveals considerable patient-to-patient variability in TNF-α pathways, correlating positively with *ERVK-7* expression. This suggests that the beneficial association between ICB response and *ERVK-7* expression may be mediated through the TNF-α pathway. Supporting our results, targeting TNF-α producing macrophages in pancreatic cancer can induce antitumor immunity(49). Moreover, our findings reveal that enhanced interferon signaling in LUAD contributes to *ERVK-7.short* overexpression. These mechanisms collectively provide insights into the potential modulation of *ERVK-7* expression in lung cancer samples.

We also found high *ERVK-7.long* expression in lung organoids derived from lung epithelial and tumor cells. However, *ERVK-7.long* expression was low in lung tissue. This difference is likely due to the composition of cell types: lung organoids consist of mostly epithelial cell types, resulting in a high proportion of cells expressing *ERVK-7.long*. In contrast, lung tissue contains a mixture of epithelial, immune, and endothelial cells, with *ERVK-7.long*-expressing cells constituting only a relatively small proportion. Coupled with lower level of chemokine signaling in normal lung compared with lung cancer, only minimal *ERVK-7.long* expression can be detected in normal lung tissue.

Moreover, we observed a slight increase in *ERVK-7.short* expression when A549 cells were treated with TNF-α (Fig. 6C), suggesting that RELA may have both direct effects on *ERVK-7.long* and indirect effects on *ERVK-7.short*. This observation can be explained by TNF-α enhancing IFN-γ receptor expression through NF-κB signaling(50), which subsequently activates IRF1 binding at the *ERVK-7.short* promoter, ultimately leading to a modest increase in *ERVK-7.short* expression following TNF-α treatment.

Finally, our scRNA-seq analysis reveals that *ERVK-7* and *ERVK-7.short* are expressed in a non-cell type specific manner. This observation suggests significant epigenetic distinctions among *ERVK-7* isoforms. Given that IFN-γ regulates *ERVK-7.short*, this isoform may play a significant role in mediating anti-tumor immune responses. A notable observation is the substantial increase in the H3K4me3 signal associated with *ERVK-7.short* in tumors (Supp. Fig. S4B). This finding contrasts with the RNA-seq data, which indicate similar expression levels of *ERVK-7.short* in both tumor and normal samples (Figure 3B). Discrepancies between RNA-seq and H3K4me3 signals have been widely reported (51–53). Further investigation is required to elucidate the transcription factors regulating *ERVK-7.short* across different cell types. Moreover, probe-based or 5’ spatial transcriptomics in normal and tumor lung tissues are crucial for providing deeper insights into the expression of *ERVK-7* transcripts across different cell types in the tumor immune microenvironment.

One aspect that remains unclear is the absence of the typical *Env* protein-coding transcript in *ERVK-7*. Further investigation is required to elucidate the mechanism by which the *Env* protein is translated from the *ERVK-7.long* and *ERVK-7.short* transcripts. Nevertheless, the discovery of TNF-α and IFN-driven promoters suggests a potential forward-feeding, inflammation-driven loop which may enhance the immunotherapy response in LUAD patients exhibiting high *ERVK-7* expression.

In summary, our discoveries highlight a specific mechanism in LUAD, where NSCLC appears to originate from SAE cells that have already acquired H3K4me3 at the *ERVK-7.long* promoter, leading to its expression and activation. The differential regulation of *ERVK-7.long* by TNF-α and IFN-γ pathways, which we found to be enriched in LUAD compared to LUSC, underscores the further upregulation of *ERVK-7* via *ERVK-7.long* in LUAD. This intricate regulation highlights *ERVK-7.long*’s potential as a biomarker and therapeutic target in NSCLC.

## Methods

### Generation of A549 Nanopore DNA-sequencing

A549 (RRID:CVCL_0023) cells were purchased from National Collection of Authenticated Cell Cultures which have been authenticated by STR analysis and mycoplasma tested prior to these experiments. The cells were seeded into a 150 mm culture dish with Ham’s F-12K (Kaighn’s) Medium and 10% fetal bovine serum. When reaching 40% confluency, cells were treated with either DMSO or 300 nM Decitabine (Abcam, ab120842) for five days. The culture medium with either DMSO or Decitabine was refreshed daily. After five days of treatment, genomic DNA was extracted from cells using the phenol/chloroform method.

### DNA methylation detection from A549 Nanopore DNA sequencing

Whole genome sequencing was performed on Nanopore PromethION 48 platform using R10.4.1 flowcells with SQK-NBD114.96 library preparation kit. A total of 45.92 Gb raw data was generated. 5mC and 5hmC modifications at CpG sites in DNA were detected and aligned against hg38 using Dorado version 0.7.0 (https://github.com/nanoporetech/dorado) with the Remora (https://github.com/nanoporetech/remora) high accuracy (HAC) model for modified basecalling (dna_r10.4.1_e8.2_400bps_hac@v5.0.0, dna_r10.4.1_e8.2_400bps_hac@v5.0.0_5mCG_5hmCG@v1). All ONT raw POD5 data were basecalled to generate modified-basecalled BAM (modBAM) files containing MM and ML tags. : SAMTOOLS, RRID:SCR_002105 (v1.20)(54) was then used to filter the modBAM files, retaining only primary alignments with mapping quality scores above 200.

### RNA-seq Alignment

For short-read, we used STAR (v2.5.3a) aligner(55) to align FASTQ files. s. For the analysis of PacBio Iso-seq data, we utilized the isoseq3 tool (https://github.com/PacificBiosciences/IsoSeq). First, *CCS* was used to compute the circular consensus sequences (CCS), *lima* and *isoseq refine* were applied to select and process CCS. Minimap2(56) was applied as an aligner to hg38. TCGA BAM files were downloaded from GDC portal (https://portal.gdc.cancer.gov/) and used with or without multimapping reads removed.

### Processing ChIP-sequencing

BWA (RRID:SCR_010910) (v0.7.18) (57) and Samtools (v1.20) (54) were used to align fastq with hg38 fasta file and create alignment BAM file. Duplicated reads were further remove using Picard (RRID:SCR_006525) (v1.128) (https://github.com/broadinstitute/picard) *MarkDuplicates*. Normalization was conducted. Chromosome X was ignored from normalization using RPGC mode in Deeptools (RRID:SCR_016366) (v3.5.4) (58) *bamCoverage*.

### scRNA-seq data processing

STAR (v2.5.3a) aligner (55) was used to align SMART-seq2 (59) FASTQ files. We used BRIE (v.2.2.2)(60) for quantification. GTF file used for BRIE is available in GitHub. Seurat (v4.4.0)(61, 62) was used to visualize, remove low-quality cells, perform dimensionality reduction and unsupervised clustering in R. For GSEA analysis, only primary LUAD tumor cells were used. Tumor cells with (*ERVK-7.long* raw reads) < 10 or 0 are considered as lowly expressing group (Low). Tumor cells with (*ERVK-7.long* raw reads) >=10 are considered as highly expressing group (High).

### Genomic sequence comparison between active LTR5Hs with LTR5Hs in ERVK-7

Among the reported active LTR5Hs elements(13), we selected those with lengths exceeding 900 bp for alignment. The coordinates of these elements are provided in Supp. Table S1. BEDTools (RRID:SCR_006646) was used to retrieve genomic sequences. Multiple sequence alignment was performed using clustalW (https://www.genome.jp/tools-bin/clustalw). Sequence difference motif image was drawn in http://weblogo.berkeley.edu/logo.cgi.

### Quantification of ERVK-7.long, ERVK-7.short and ERVK-7

All bulk RNA-seq data, including datasets from GEO and TCGA, were quantified using featureCounts (RRID:SCR_012919) (v2.0.1)(63). We created a custom GTF file by combining Gencode basic v44(12) and the *ERVK-7* GTF file (green track, Fig. 3A). TCGA data was analyzed using the Cancer Genomics Cloud(64). We separately quantified the region (trans996c17b9912b02e8), where the *ERVK-7* envelope protein is known to be coded using featureCounts. All of the GTF files are uploaded on GitHub.

### TCGA Downstream analysis

A two tailed t-test was performed to compare TPM-normalized values of *ERVK-7* transcripts across LUAD tumor, LUSC tumor, and normal samples. These samples include both male and female subjects where randomization and blinding were performed in accordance to TCGA. Spearman’s rank correlation coefficient was used to compared with expression of different *ERVK-7* transcripts. TCGA GSEA analysis was conducted with clusterProfiler (v3.18.1)(65, 66). The TNF-α signaling via NF-κB pathway and interferon-γ signaling gene sets were obtained from the MSigDB Hallmark collection(67). The TNF receptor pathway gene set was sourced from the NCI-Pathway Interaction Database (NCI-PID)(68). A multivariable regression model was applied using the following variables: amplification state of 1q22 (defined as “amplified” for copy number ≥ 3, and “none” for others), and single-sample Gene Set Enrichment Analysis (ssGSEA) scores(69) for TNF receptor and IFN-γ signaling pathways. All values were z-score normalized prior to analysis.

### Quantification of ERVK-7 transcripts contributions in TCGA

Assuming *ERVK-7* expression is derived solely from *ERVK-7.long* and *ERVK-7.short*:

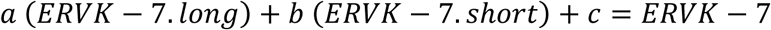

We first utilized a multivariate regression model with *ERVK-7.long* and *ERVK-7.short* as variables to obtain coefficients for each transcript: *a* for *ERVK-7.long* and *b* for *ERVK-7.short*. All values were Z-scored prior to applying the regression model. We then multiplied the coefficients as normalization factors to TPM values for each patient. Finally, we calculated log2 (normalized *ERVK-7.long* / normalized *ERVK-7.short*) to quantify the relative contribution of the transcripts to *ERVK-7* expression.

### Visualization and downstream

We utilized UCSC Genome Browser (RRID:SCR_005780) (https://genome.ucsc.edu) and IGV (70) for visualization of tracks. All of the plots were generated in R studio. Diagrams were refined with Biorender (RRID:SCR_018361). All of the RNA-seq values were normalized using TPM with library size of genes.

## Data and Code Availability

Data from Direct WGS of A549 generated using Oxford Nanopore Technologies has been deposited in SRA under PRJNA1178369. Repeatmasker annotations was downloaded from UCSC table browser hg38 (https://genome.ucsc.edu/). Three histone ChIP-seq (H3K4me1, H3K4me3, H3K27ac), WGBS for A549 and normal lung tissue, and three histone ChIP-seq (H3K4me1, H3K4me3, H3K27ac) for three primary cells were retrieved from ENCODE Project (https://www.encodeproject.org/) with accession code listed in Supp. Table S4. TCGA BAM files are transferred from GDC portal (https://portal.gdc.cancer.gov/ ) to CGC (https://www.cancergenomicscloud.org/) for downstream analysis. SMART-seq2 lung scRNA cohort study is available via ENA PRJNA591860 (https://www.ebi.ac.uk/ena/browser/view/PRJNA591860). Others are retrieved from GEO (https://www.ncbi.nlm.nih.gov/geo/) with the accession codes listed in Supp. Table S4. Code and GTF files used in the analysis and figure generation are available on GitHub (https://github.com/jasonwong-lab/ERVK-7.git)

Funding

This work was supported by the Centre for Oncology and Immunology under the Health@InnoHK Initiative funded by the Innovation and Technology Commission, the Government of Hong Kong SAR, China, and the National Key R&D Program of China (2023YFC2508901).

