## Supplemental Figures for "A tumor necrosis factor-α responsive cryptic promoter drives overexpression of the human endogenous retrovirus *ERVK-7*"

**Supporing information**

| **Table of contents** | Page |
|---|---|
| Figure S1 | S-1 |
| Figure S2 | S-2 |
| Figure S3 | S-3 |
| Figure S4 | S-4 |
| Figure S5 | S-5 |
| Figure S6 | S-6 |
| Figure S7 | S-7 |
| Figure S8 | S-8 |
| Figure S9 | S-9 |
| Figure S10 | S-10 |
| Table S1 | Separate Excel file S-1 |
| Table S2 | Separate Excel file S-2 |
| Table S3 | Separate Excel file S-3 |
| Table S4 | Separate Excel file S-4 |

### Supplementary Figures

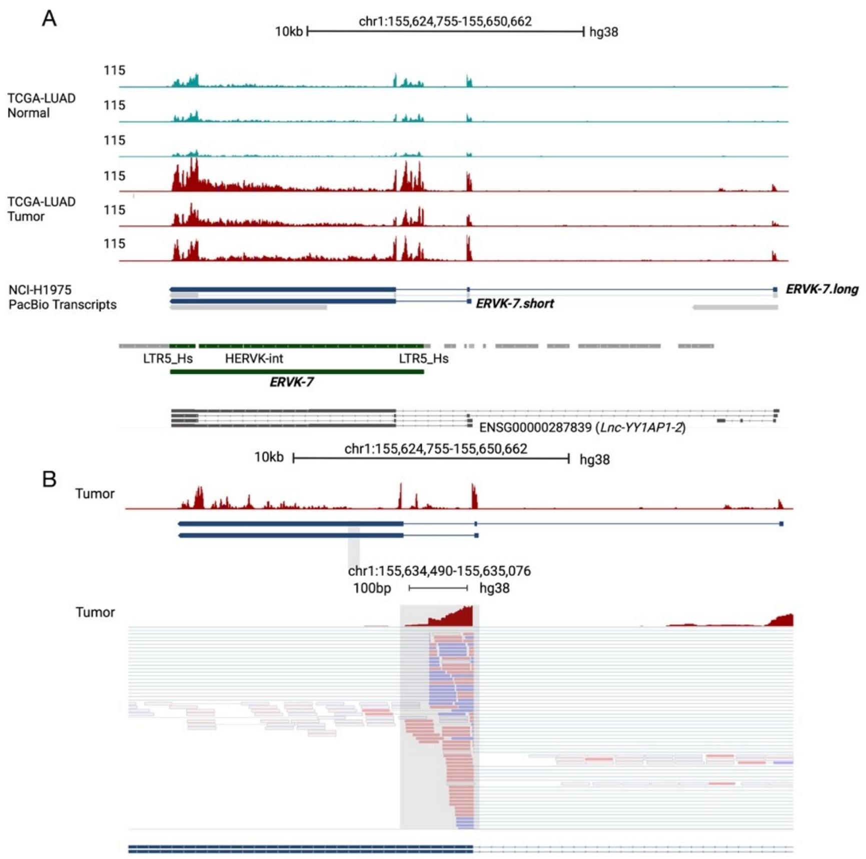


**Supp. Figure S1**

(A) First six tracks show three pairs of TCGA LUAD normal (Blue) and tumor (Red) patients (TCGA-44-2668, TCGA-38-4632, TCGA-44-6147) including both unique and multi-mapped reads coverage. The below track shows the NCI-H1975 (LUAD cell line) Pacbio Iso-Seq transcript. The first blue-colored transcript and the second blue-colored transcript represent *ERVK-7.long* and *ERVK-7.short* respectively. The track below represents RepeatMasker annotation. Regions corresponding to *ERVK-7* are colored green. The final track (Grey) shows Gencode V46 comprehensive. All of the transcripts are named long noncoding RNA for *YY1AP1*. (B) Top: TCGA-44-3396 tumor sample. Bottom: Zoomed in the region for the grey area in the top box. Demonstrating reads going inside the *ERVK-7* internal part.

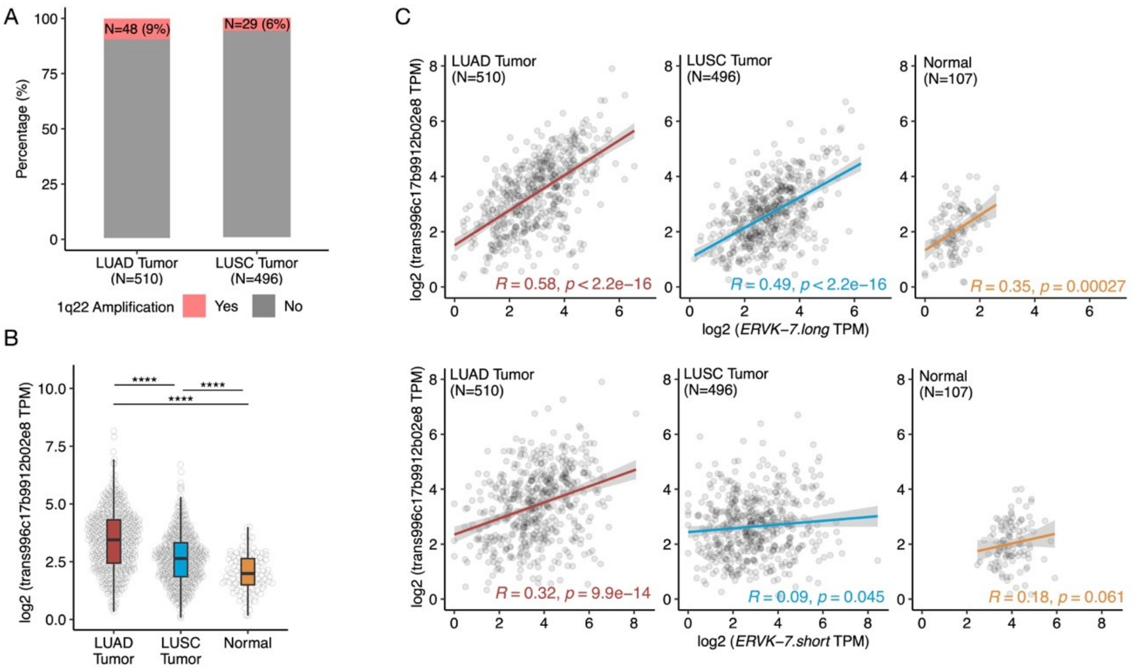


**Supp. Figure S2**

(A) Expression of trans996c17b9912b02e8, from which *ERVK-7 Env* protein is encoded. Log2 TPM values used. (NS: p > 0.05, *: p ≤ 0.05, **: p ≤ 0.01, ***: p ≤ 0.001, ****: p ≤ 0.0001). (B) Correlation plots with *ERVK-7 Env* protein coding region. Log2 TPM values used. *ERVK-7.long* (top) and *ERVK-7.short* (bottom).

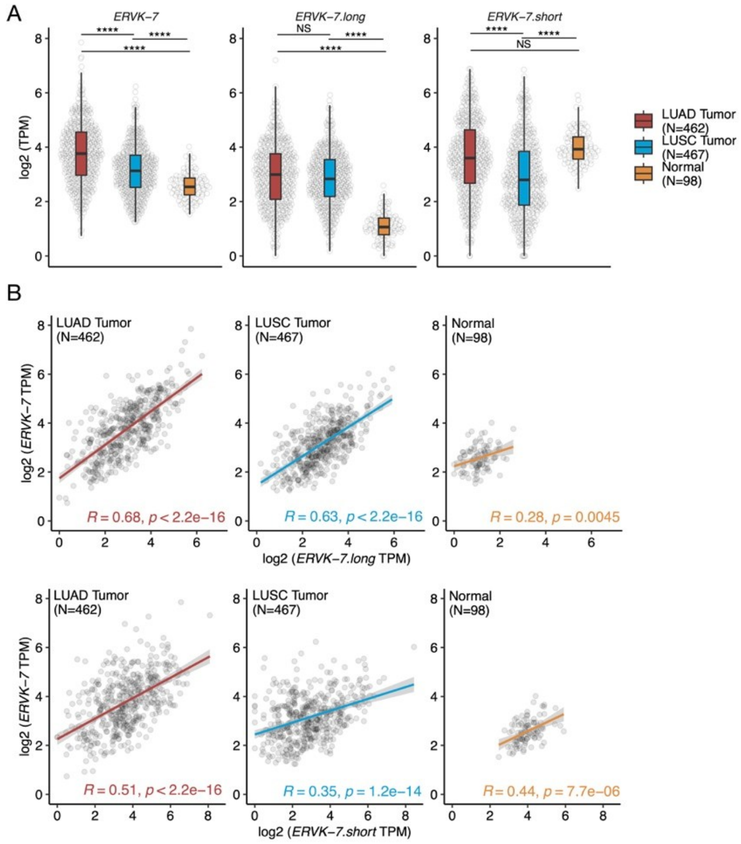


**Supp. Figure S3**

(A) Expression of *ERVK-7* in patients without amplification og 1q22. Log2 TPM values used. (NS: $p > 0.05$, *: $p \le 0.05$, **: $p \le 0.01$, ***: $p \le 0.001$, ****: $p \le 0.0001$). (B) Correlation plots with *ERVK-7*. Log2 TPM values used. *ERVK-7.long* (top) and *ERVK-7.short* (bottom).

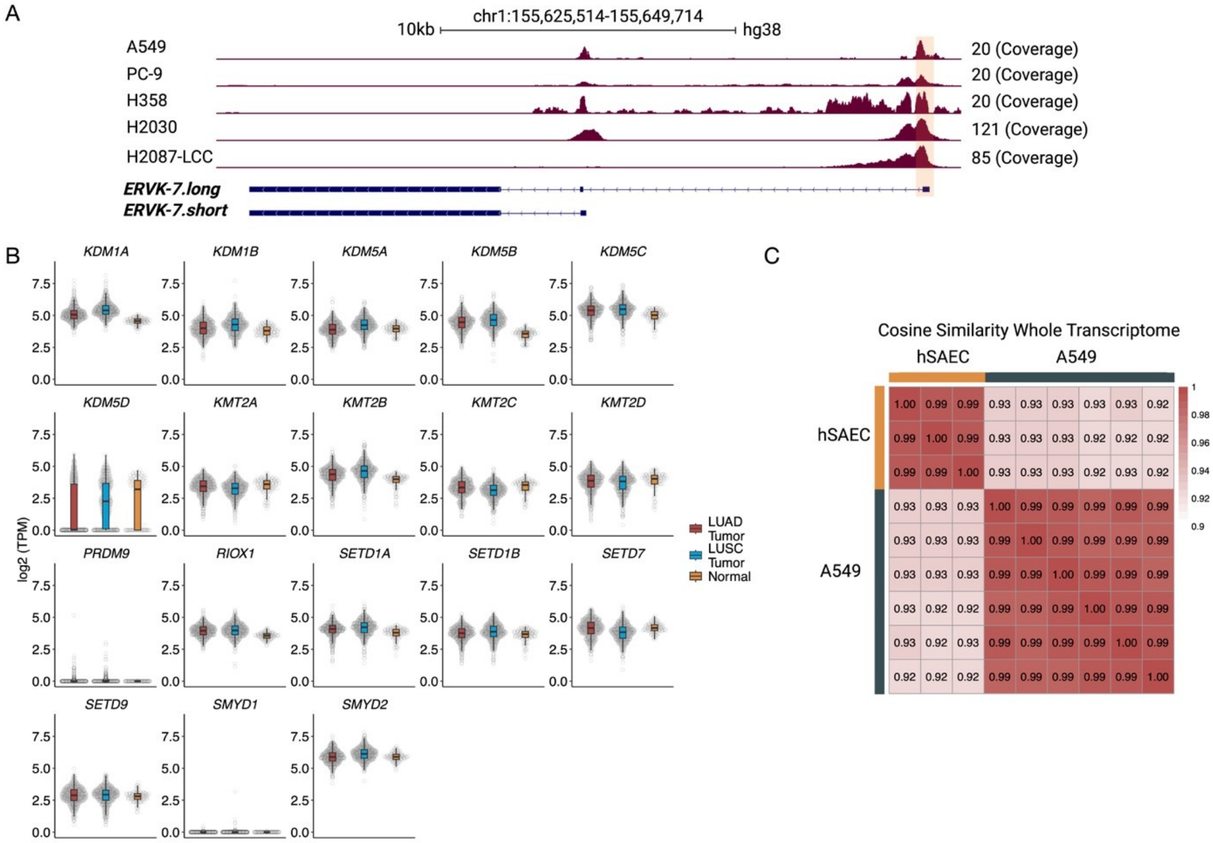


**Supp. Figure S4**

(A) *ERVK-7* region H3K4me3 coverage for five NSCLC cell lines. (B) Known H3K4me1 or H3K4me3 writer and eraser protein-coded genes. Log2 TPM used. LUAD tumor (Red), LUSC tumor (Blue), LUAD and LUSC normal (Yellow). (C) Cosine similarity between hSAEC cell line and A549 cell line for the whole transcriptome. The value/color represents the cosine similarity value.

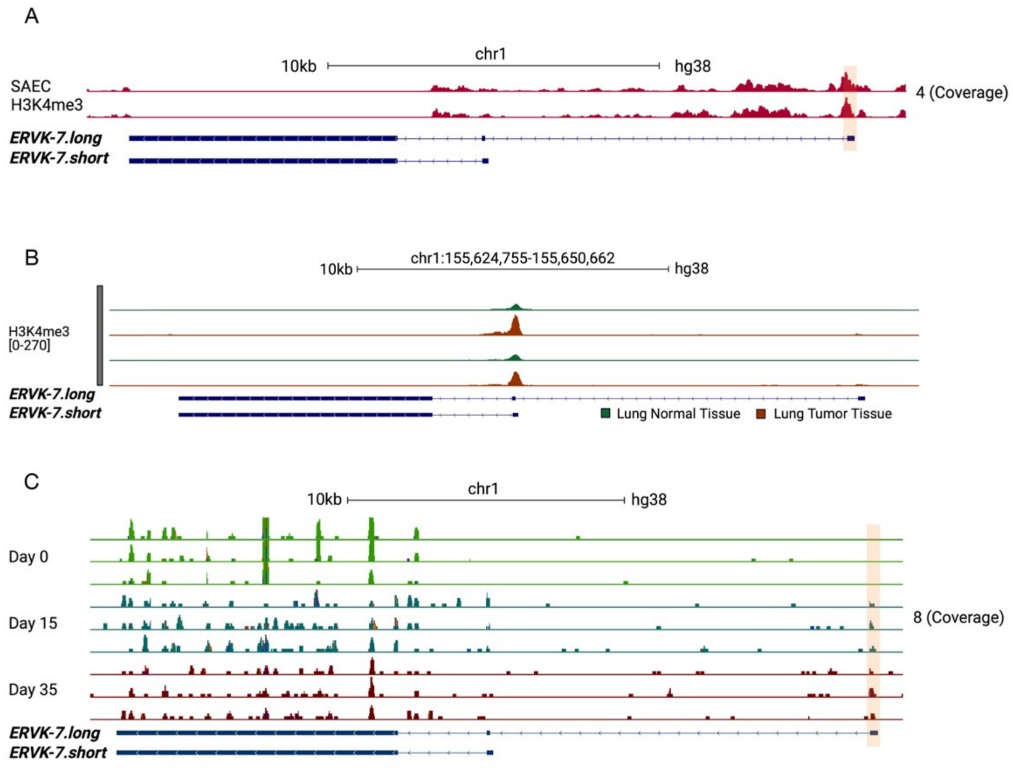


**Supp. Figure S5**

(A) Two replicates of H3K4me3 ChIP-seq for hSAEC samples. Orange area highlights the promoter of *ERVK-7.long* transcript. (B) Two datasets of H3K4me3 ChIP-seq compare normal lung (green) and lung tumor (red) tissues. (C) Three replicates of human pluripotent stem cell (PSCs) at day 0 (Green), day 15 (Blue), day 35 (Red). Orange area highlights the promoter of *ERVK-7.long* transcript.

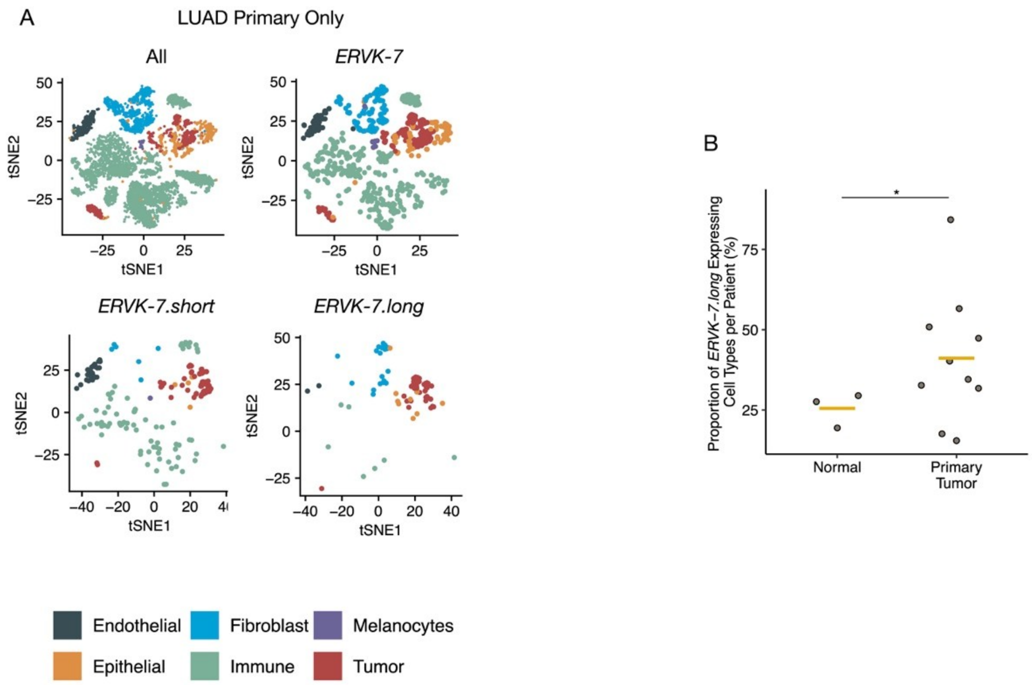


**Supp. Figure S6**

(A) T-distributed stochastic neighbor embedding (t-SNE) plot of SMART-seq2 scRNA-seq data from lung cancer and normal samples. Subpanels show all cells or cells expressing specific *ERVK-7* transcripts. Six board cell types marked: Dark grey: Endothelial, Blue: Fibroblast, Purple: Melanocytes, Orange: Epithelial, Green: Immune, Red: Tumor. (B) Proportion of *ERVK-7.long* expressing cell types per patients. Yellow line indicates mean of each group. Student t-test performed. (*: $p \leq 0.05$).

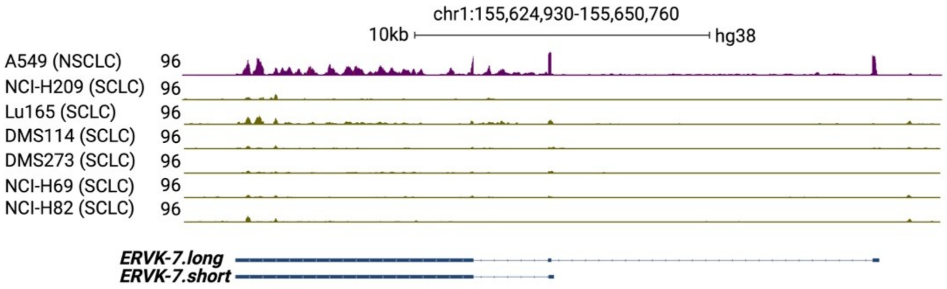


**Supp. Figure S7**

*ERVK-7* region RNA-seq read coverage for A549 (Purple) and 6 SCLC samples (Green).

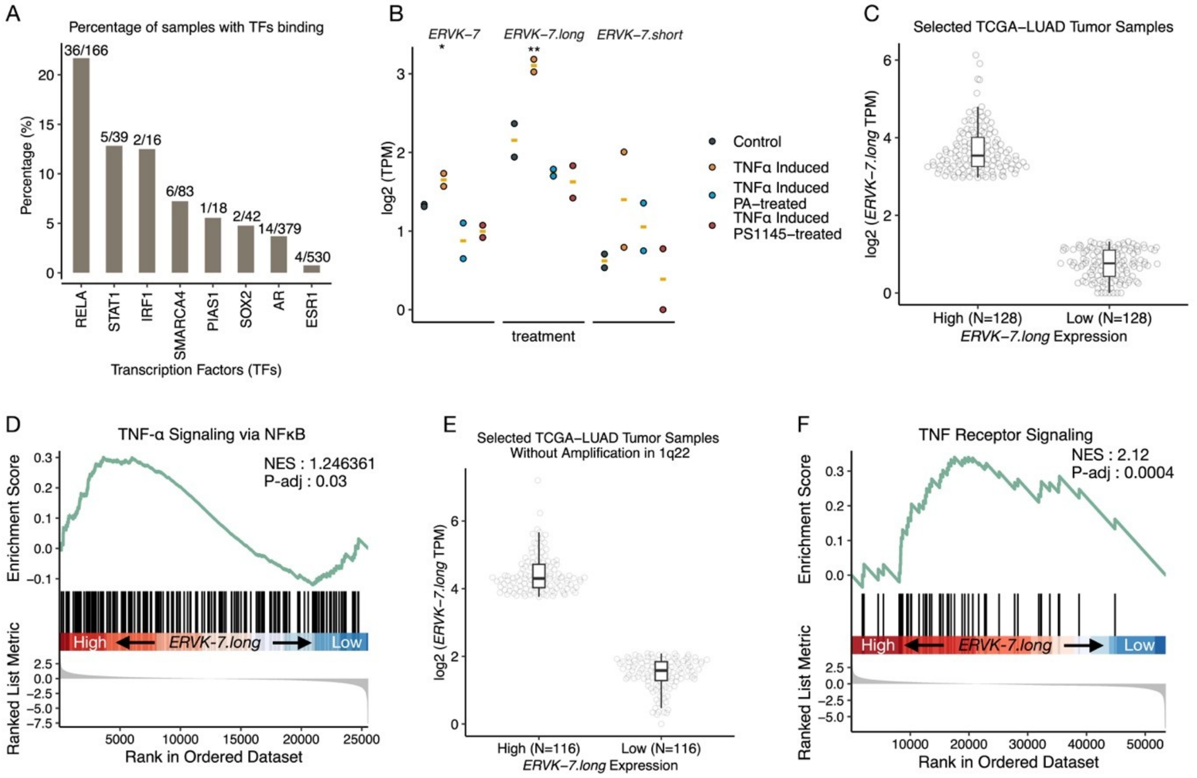


**Supp. Figure S8**

(A) Percentage of transcription factor (TF) binding at the promoter of *ERVK-7.long* based on data from CistromeDB. For each TF, the percentage of samples binding over the total is calculated and represented above each bar. (B) log2 TPM gene expression values for three *ERVK-7* related transcripts. PA and PS1145 are inhibitors for TNF-α. A549 (Black), TNF-α (Orange), TNF-α with PA-treated (Blue), and TNF-α with PA-treated (Red). Log2 TPM used. ANOVA test conducted. (NS: $p > 0.05$, *: $p \leq 0.05$, **: $p \leq 0.01$, ***: $p \leq 0.001$, ****: $p \leq 0.0001$). (C) *ERVK-7.long* expression from TCGA-LUAD tumor samples based on top 25% of patients with high *ERVK-7.long and* the bottom 25% of patients with low *ERVK-7.long* that were selected for GSEA analysis. (D) GSEA plot for TNF-α signaling via NFκB pathway between TCGA-LUAD tumor samples high vs low groups in (C). (E) *ERVK-7.long* expression in TCGA-LUAD tumor samples without 1q22 amplification, comparing the top 25% of patients with high *ERVK-7.long* expression to the bottom 25% with low expression selected for GSEA analysis. (F) GSEA plot for TNF receptor signaling between TCGA-LUAD tumor samples high vs low groups in (E).

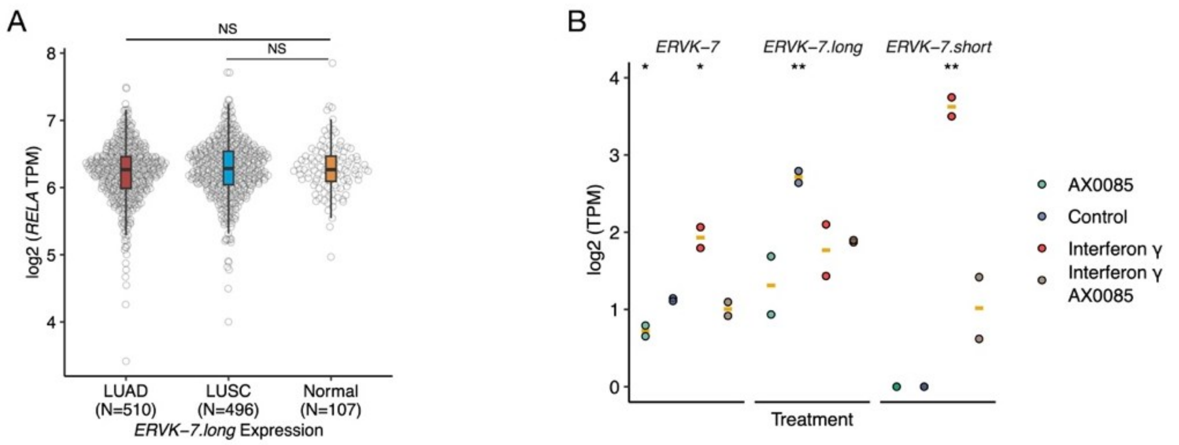


**Supp. Figure S9**

(A) Log2 TPM values for *RELA* in LUAD tumor (Red), LUSC tumor (Blue), and Normal lung (Yellow). Student t-test performed. (B) Log2 TPM values for three *ERVK-7*-related transcripts. AX0085 is an inhibitor of IFN-γ. A549 (Blue), IFN-γ (Red), AX0085 (Green), and IFN-γ with AX0085 (Brown). Log2 TPM used. ANOVA test conducted. (NS: $p > 0.05$, *: $p \leq 0.05$, **: $p \leq 0.01$, ***: $p \leq 0.001$, ****: $p \leq 0.0001$).

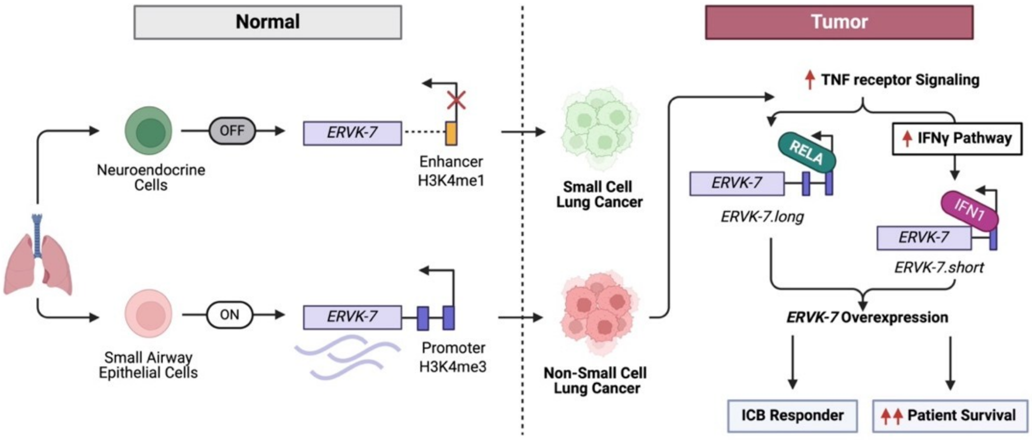


**Supp. Figure S10**

Diagram Illustrating General Regulation of *ERVK-7* transcripts in Lung Cancer
